# Bone marrow-derived macrophages loaded with boron carbide nanoparticles as selective boron carriers for crossing the blood-brain barrier toward the glioma microenvironment for boron neutron capture therapy

**DOI:** 10.64898/2026.01.09.698353

**Authors:** Anna Rudawska, Agnieszka Szczygieł, Katarzyna Węgierek-Ciura, Jagoda Mierzejewska, Dawid Kozień, Paulina Żeliszewska, Monika Chaszczewska-Markowska, Piotr Rusiniak, Katarzyna Wątor, Zbigniew Pędzich, Elżbieta Pajtasz-Piasecka, Bożena Szermer-Olearnik

## Abstract

**Background:** Boron neutron capture therapy (BNCT) is a type of targeted radiotherapy that destroys boron-containing cancer cells using a neutron beam. It is intended for the treatment of patients with tumors that are resistant to conventional treatment, inoperable and recurrent. Therefore, it represents a promising therapy for the treatment of glioblastoma multiforme, the most aggressive stage IV cancer. To ensure high efficacy of BNCT, it is necessary to use selective boron carriers that deliver therapeutic doses of the boron-10 isotope to the tumor site, demonstrate a lack of systemic toxicity, and, most importantly, cross the blood-brain barrier. In our studies, we propose using bone marrow-derived macrophages as cellular carriers of boron carbide nanoparticles to selectively deliver boron-10 to the tumor microenvironment. Macrophages can efficiently engulf foreign particles, migrate and accumulate at the tumor site, as well as cross the blood-brain barrier, making them extremely promising boron carriers for BNCT.

**Results:** Bone marrow-derived macrophages in three polarization states (M0, M1, and M2) demonstrated a high uptake ability of boron carbide nanoparticles at a concentration of 100 µg/ml during 24-hour incubation. The highest boron concentration was detected in M1 macrophages, resulting in the greatest inhibition of viability and slower spontaneous migration of this population compared to other types of macrophages. Nevertheless, all macrophage populations (M0, M1, and M2) loaded with boron carbide nanoparticles migrated comparably through the brain endothelial cell layer, mimicking the blood-brain barrier, toward the CCL2-rich supernatant of GL-261 glioma cells. Furthermore, macrophages with nanoparticles did not affect the viability of glioma spheroids after 6 days of co-culture, in contrast to macrophages without nanoparticles, which increased the survival of tumor cells. Importantly, M1 macrophages did not repolarize toward the M2 phenotype during co-culture, as evidenced by the stable expression of CD206 in these cells. In addition, M2 macrophages loaded with B_4_C nanoparticles showed lower CD206 expression after contact with spheroids compared to this macrophage population without nanoparticles.

**Conclusions:** Bone marrow-derived macrophages loaded with boron carbide nanoparticles showed great potential for use in BNCT as boron carriers crossing the blood-brain barrier toward the glioma microenvironment.

## Background

Glioma is the most common primary malignant brain tumor in adults, accounting for approximately half of all primary tumors of the central nervous system [1]. It originates from the glial cells of the brain, and its histological classification is based on four grades of malignancy (I-IV). Glioblastoma multiforme is the most prevalent and aggressive stage IV tumor with a poor prognosis. Standard treatment includes surgical resection supplemented with radiotherapy and temozolamide chemotherapy. However, due to treatment resistance, conventional therapies are not effective for many patients, resulting in frequent tumor recurrence and high mortality [2]. Effective treatment of gliomas remains a significant challenge due to the tumor’s location in hard-to-reach areas of a crucial organ, its heterogeneous nature, the immunosuppressive tumor microenvironment (TME), and the presence of the blood-brain barrier (BBB), which is impenetrable to many drugs [3].

The high resistance of gliomas to conventional treatments increases the need for new effective therapies. One promising treatment strategy is boron neutron capture therapy (BNCT), which is a type of targeted radiotherapy that destroys tumors at the cellular level. BNCT was developed to treat patients with unresectable, recurrent, and locally advanced cancers. Since its first clinical trials in the 1950s, it has been used to treat patients with malignant gliomas. However, BNCT is also utilized in the treatment of melanomas, as well as recurrent head and neck cancers [4]. The basis of BNCT is irradiation of the tumor containing the accumulated boron-10 (^10^B) isotope with thermal or epithermal neutrons. The reaction results in the emission of an alpha particle (^4^He) and a recoiling lithium-7 nucleus (^7^Li) with high linear energy transfer (LET). The released energy is deposited within a single tumor cell radius (<10 μm), limiting the destructive effect on normal cells [5].

Despite many advantages, BNCT remained in the clinical trial phase for years. In 2020, Japan was the first country to approve BNCT for hospital use and provide health insurance coverage for the treatment [6]. The reason for the lack of widespread implementation of this therapy is the deficiency of boron carriers that selectively accumulate in cancer cells, providing the required boron concentrations (>20 μg ^10^B per gram of tumor or > 10^9^ atoms per cell). Furthermore, boron carriers should be characterized by a lack of systemic toxicity, maintaining a constant boron concentration during neutron irradiation, as well as rapid clearance from the blood and normal tissues. Additionally, in the case of brain tumors, the ability to cross the BBB is necessary. Therefore, there is a great need to find a boron carrier that meets all the requirements [7].

One of the promising candidates for BNCT is boron carbide (B_4_C), characterized by high neutron absorptivity. Moreover, B_4_C can form diverse stoichiometries with a high boron content [8]. In our previous work, we confirmed that originally synthesized boron carbide nanoparticles delivered a high concentration of boron (~7.8 mg/l of boron per million cells) to cancer cells [9]. However, to increase the selectivity of delivery of this compound to the tumor microenvironment and reduce undesirable interactions with normal cells, the use of targeted delivery methods is crucial. For this purpose, we propose using a “Trojan horse” strategy based on macrophages as cellular carriers of boron carbide nanoparticles. Macrophages exhibit natural phagocytic abilities to engulf foreign particles, as well as low immunogenicity, biocompatibility, and long viability. Importantly, macrophages cross biological barriers, including the BBB, and due to their tropism toward hypoxia, they efficiently migrate and accumulate in the tumor microenvironment [10]. In gliomas, tumor-associated macrophages (TAMs) forming the TME may constitute 30-50% of the tumor mass. TAMs found in gliomas originate from microglia present in the brain, but also from bone marrow-derived macrophages (BMDMs) [11]. The main factor responsible for the migration and recruitment of monocytes and macrophages from the blood circulation to the brain tumor is C-C motif chemokine ligand 2 (CCL2), also known as monocyte chemotactic protein-1 (MCP-1). CCL2, abundantly produced by glioma cells, binds to the C-C chemokine receptor type 2 (CCR2) present on monocytes/macrophages and promotes their differentiation into TAMs [12].

Bone marrow-derived macrophages present in TME possess diverse phenotypes and functions that are acquired in response to environmental stimuli due to their phenotypic plasticity. The main classification of macrophages includes classically activated macrophages (M1) and alternatively activated macrophages (M2), among which the following subpopulations are distinguished: M2a, M2b, M2c, and M2d [13]. M1 cells produce proinflammatory cytokines and chemokines such as IL-1α, IL-1β, IL-6, IL-12, and TNF-α, stimulating Th1 lymphocytes and participating in antitumor responses and inhibiting tumor progression. M1 macrophages present antigens via the major histocompatibility complex class II (MHC II) and also express the costimulatory molecules CD80 and CD86 on their surface [14]. M2 macrophages secrete anti-inflammatory cytokines (IL-4, IL-10, and IL-13), chemotactic and angiogenic factors, as well as extracellular matrix components, promoting tumor progression. M2 cells, creating an immunosuppressive TME, inhibit the action of cytotoxic T lymphocytes and attract regulatory T lymphocytes (Tregs) and myeloid-derived suppressor cells (MDSCs). Furthermore, M2 cells overexpress mannose receptors such as CD206 and CD209. Due to the immunosuppressive effect of the tumor microenvironment, M1 macrophages may undergo repolarization to the M2 phenotype, resulting in the predominance of M2 macrophages among TAMs [11, 15].

In our previous studies, we demonstrated a relationship between the polarization state of bone marrow-derived macrophages (M0, M1, and M2) and their sensitivity to boron carbide nanoparticles [16]. We also confirmed the ability of all BMDM populations to effectively engulf and accumulate B_4_C nanoparticles. To complement and extend these studies, the current work focuses on the crucial aspects of using BMDMs as boron carriers in BNCT. First, we present confirmation of the efficient engulfment of nanoparticles by macrophages based on analysis of cellular boron concentration. Furthermore, we compare the migratory capacity of all BMDM populations loaded with B_4_C nanoparticles, both spontaneously and through a brain endothelial cell layer mimicking the BBB, toward the CCL2-rich supernatant of GL-261 glioma cells. In the final stage of our research, we analyze the effect of macrophages on the viability of spheroids obtained from glioma cells and assess changes in the macrophage phenotype after co-culture with the spheroids.

## Materials and Methods

### Boron carbide nanoparticles

Boron carbide (B_4_C) nanoparticles used in this study were synthesized by the vapor deposition and had a size of 32 ± 10 nm. Detailed characterization of the obtained B_4_C nanoparticles was performed using dynamic light scattering (DLS), laser Doppler velocimetry (LDV), atomic force microscopy (AFM), X-ray diffraction analysis (XRD), scanning electron microscope (SEM), and transmission electron microscope (TEM), as described in the work of Kozień et al. [17].

### Cell culture

Mouse bEnd.3 endothelial cells obtained from the American Type Culture Collection (ATCC; CRL-2299) were maintained in Dulbecco’s modified Eagle’s medium (DMEM; ATCC) containing 10% heat-inactivated fetal bovine serum (FBS; ATCC), 100 U/ml penicillin, and 100 mg/ml streptomycin (all from Sigma-Aldrich).

Mouse glioma GL-261 cells obtained from the Leibniz Institute DSMZ (German Collection of Microorganisms and Cell Cultures GmbH; ACC 802) were maintained in DMEM (ATCC) supplemented with 10% FBS, 100 U/ml penicillin, and 100 mg/ml streptomycin (all from Sigma-Aldrich).

All cell cultures were maintained in a NUAIRE CO_2_ incubator (37°C, 5% CO_2_, 95% humidity).

### Preparation of bone marrow-derived macrophages

Femurs and tibias were collected from healthy 7-8-week-old female C57BL/6 mice. To isolate bone marrow cells, the marrow cavities of the cleaned bones were flushed with RPMI-1640 medium (Gibco) supplemented with 3% FBS, 100 U/ml penicillin, and 100 mg/ml streptomycin (all from Sigma-Aldrich). Isolated bone marrow cells were centrifuged for 7 min at 192 × g at 4°C, and placed on T75 culture flasks (Corning). Cells were cultured in RPMI-1640 medium (Gibco) with the addition of 1 mM sodium pyruvate, 0.05 mM 2-mercaptoethanol, 10% FBS, 100 U/ml penicillin, 100 mg/ml streptomycin (all from Sigma-Aldrich), and 50 ng/ml recombinant mouse macrophage colony-stimulating factor (M-CSF; ImmunoTools), called culture medium. The medium was replaced with a fresh one every 2 days. After 8 days of culture under these conditions, unpolarized (M0) bone marrow-derived macrophages were obtained. To activate macrophages, after 7 days of culture with M-CSF, 20 ng/ml interferon-γ (IFN-γ; ImmunoTools) and 100 ng/ml lipopolysaccharide (LPS, from E. coli O111:B4; Sigma Aldrich) were added for 24 h to polarize to M1 macrophages, and 20 ng/ml interleukin 4 (IL-4; ImmunoTools) to polarize to M2 macrophages.

### Phenotypic characterization of BMDMs by flow cytometry

Phenotypic characterization was performed using flow cytometry to confirm the differentiation of bone marrow cells toward macrophages, their activation status, and the lack of influence of B_4_C nanoparticles on antigen expression. Briefly, BMDMs were stained with cocktails of fluorochrome-conjugated monoclonal antibodies: anti-F4/80 Alexa Fluor 700, anti-CD11b PerCP-Cy5.5, anti-MHC II FITC, anti-CD86 PE-Cy7 (all from BioLegend), anti-CD40 PE (Becton Dickinson), and appropriate isotype controls. Then, the cells were fixed using the Foxp3/Transcription Factor Staining Buffer Set (Thermo Fisher Scientific) and incubated with anti-CD206 APC (BioLegend) antibody. Analysis was performed using an LSRFortessa flow cytometer with Diva Software (Becton Dickinson). The graphs and histograms were prepared in GraphPad Prism 10 (GraphPad Software) and NovoExpress 1.3.0 software (ACEA Biosciences, Inc.). A detailed scheme of BMDM analysis was published in the work of Wróblewska et al. [16]

Additionally, to evaluate the influence of glioma spheroids on changes in the expression of BMDM-specific antigens, phenotypic characterization was performed on days 3 and 6 of co-culture as described above. The only exception to the procedure was the separation of spheroids into single cells using the LifeGel Digestion kit (RealResearch) before staining with the antibody cocktail. The phenotypic characterization of BMDMs after 3 and 6 days of co-culture with glioma spheroids was performed in two independent experiments.

### Formation of glioma spheroids

GL-261 cells were seeded at a density of 2 × 10^3^ cells/well in 96-well plates coated with LifeGel (Real Research) in DMEM (ATCC) containing B-27 supplement (Gibco), 20 ng/ml epidermal growth factor (EGF; ImmunoTools), 20 ng/ml fibroblast growth factor (FGF; ImmunoTools), 100 U/ml penicillin, and 100 mg/ml streptomycin (Sigma-Aldrich). From day 6 of culture, the culture medium was replaced every 2 days to ensure spheroid growth.

### Co-culture of glioma spheroids and BMDMs

On the 8^th^ day of culture, M0, M1, and M2 macrophages, both with or without B_4_C nanoparticles, were placed onto 10-day-old GL-261 spheroids at a density of 5 × 10^3^ cells/well in RPMI-1640 medium (Gibco) supplemented with 10% FBS, 100 U/ml penicillin, and 100 mg/ml streptomycin (all from Sigma-Aldrich). Co-culture was conducted for 3 and 6 days at 37°C (5% CO_2_, 95% humidity).

For fluorescence microscopy imaging, macrophages were stained with 2 µM Calcein AM Green (Thermo Fisher Scientific) before addition to spheroids. GL-261 spheroids with and without BMDMs were observed using an Olympus IX81 inverted fluorescence microscope.

### MTT cell viability assay

M0, M1, and M2 bone marrow-derived macrophages were placed in 96-well plates at a density of 2 × 10^4^ cells/well. After one day, boron carbide nanoparticles were added to a final concentration ranging from 0.1 to 400 μg/ml and incubated for 24 h. MTT dye (3-(4,5-dimethylthiazol-2-yl)-2,5-diphenyltetrazolium bromide; Sigma-Aldrich) was then added at 5 mg/ml for 4 h, followed by lysing overnight in lysis buffer (N,N-dimethylmethanamide, sodium dodecyl sulfate, and water) at 37°C. The absorbance of the dissolved formazan crystals was measured at 570 nm using a BioTek Epoch 2 plate reader with Gen5 Software (Agilent Technologies). MTT assays were performed in two independent experiments.

### Holotomography

M0, M1, and M2 bone marrow-derived macrophages were seeded on a TomoDish (Altium) at a density of 2 × 10^5^ cells in 2 ml of culture medium. After attaching the cells to the dish, B_4_C nanoparticles were added to a final concentration of 100 μg/ml for 24 h. Afterwards, the culture medium containing noninternalized nanoparticles was removed and replaced with fresh medium. The cells were visualized with an HT-2H commercial optical diffraction tomography microscope (Tomocube Inc). Three-dimensional refractive index (RI) tomograms were created from a series of two-dimensional holographic images obtained from 49 illumination conditions. The diffracted beams from the samples were collected using a high numerical aperture (NA = 1.2) objective lens UPLSAP 60XW (Olympus). Data processing and visualization were performed with TomoStudio HT-2H Gen3-3.3.9 software (Tomocube Inc).

### Analysis of the boron content in BMDMs using ICP-MS

#### Preparation of samples

M0, M1, and M2 bone marrow-derived macrophages were placed in 24-well plates at a density of 2 × 10^5^ cells/well. Next, boron carbide nanoparticles were added to a final concentration of 100 μg/ml and incubated for 24 h. After this time, the medium above the cells was removed, and the cells were washed three times with phosphate-buffered saline (PBS). Proteinase K (Sigma-Aldrich) solution at a concentration of 1 mg/ml in 100 mM sodium bicarbonate (POCH) was then added and incubated for 30 min at 50°C. Cells were harvested from plates, and the wells were washed 2 times with MiliQ water to collect all cells.

#### Microwave digestion of samples

The samples under investigation had to be dissolved in a homogeneous solution before instrumental analysis. This process uses the modern ultraWAVE mineralization system (Milestone Srl). First, the material was well-mixed using a vortex mixer. Approximately 0.5 g of each material was sampled and placed into Teflon vessels, followed by the addition of 5 ml of concentrated 67% ultrapure nitric acid (V) (NORMATOM, Ultrapure for trace metal analysis). The vessels were subsequently positioned in a microwave mineralizer. The mineralization procedure involved heating to 250°C over 25 min and maintaining the target temperature for 20 min, with the microwave power set to 1500 W. After digestion, the resulting solutions were quantitatively transferred to 10 ml polypropylene vessels (borosilicate glassware was avoided to prevent boron contamination) and diluted to a final mass of 10 g with deionized water obtained from the Direct-Q 3 UV system (Merck Millipore). The prepared solutions were then subjected to instrumental analysis to determine the total boron concentration in the samples.

#### Instrumental analysis of the boron content by ICP-MS

After microwave digestion, samples containing bone marrow-derived macrophages both with and without boron carbide nanoparticles were analyzed using the inductively coupled plasma mass spectrometry (ICP-MS) technique on the iCAP RQ (C2) instrument (Thermo Fisher Scientific) in accordance with the ISO 17294–2 standard [18]. A five-point calibration curve, including a blank, was prepared for the analysis, covering concentrations ranging from 5 to 500 μg/l. Both the certified boron reference materials applied for calibration and the laboratory control samples used to verify the calibration curve were produced in accordance with ISO 17034 standards [19]. The certified reference materials used were Boron B - 10 g/l in H_2_O for ICP (CPAchem Ltd) and ICP multielement standard solution IV (Supelco Analytical Products). Quantitative determination was carried out using the isotope ^10^B, which occurs naturally at about 20% abundance. Based on the measurements, the estimated limit of quantification was 1 μg/l, while the uncertainty associated with boron determination by ICP-MS was 16%. Final results were corrected for the sample mass subjected to mineralization and the diluted final mass. Measurements were performed on three independent biological samples for each macrophage population. The obtained boron concentration in the samples was converted to million cells.

### Scratch assay

M0, M1, and M2 bone marrow-derived macrophages were placed at a density of 8 × 10^5^ cells/well in 6-well plates in the culture medium. B_4_C nanoparticles at a final concentration of 100 μg/ ml were added for 24-hour incubation. Next, a gentle scratch was made with a 200 μl tip in the center of the cell monolayer in the wells. The floating cells were removed by washing with PBS, and then the culture medium was added. The macrophages were incubated at 37°C (5% CO_2_, 95% humidity) and imaged over time for 24 h using an Olympus CKX53 light microscope. Two independent experiments were performed. The images show a selected part of the well for each sample at 0, 4, and 24 h.

### Determination of CCL2 production

Culture supernatant of adherent GL-261 cells was collected after 24, 48, and 72 h. The supernatant was also collected after 3 and 6 days of co-culture of GL-261 spheroids and BMDMs. The concentration of CCL2 was determined using a commercially available ELISA kit (Invitrogen). The test was conducted according to the manufacturer’s instructions, with each sample tested in triplicate.

### Transwell migration assay

To generate a monolayer of brain endothelial cells imitating the blood-brain barrier, 7.5 × 10^4^ bEnd.3 cells were placed on Transwell inserts (pore size: 3 μm; Corning) in 150 µl of DMEM supplemented with 10% FBS, 100 U/ml penicillin, and 100 mg/ml streptomycin per insert, and placed in a 24-well plate. After 3 days, the culture medium on the inserts was replaced with fresh medium. On day 5 of bEnd.3 cell culture on the inserts, when a tight monolayer was achieved, the Transwell migration assay was initiated.

M0, M1, and M2 macrophages were incubated with boron carbide nanoparticles at a concentration of 100 μg/ml for 24 h. Next, all macrophage populations, both with and without B_4_C nanoparticles at a density of 5 × 10^4^ cells/insert, were applied to the bEnd.3 cell-coated inserts in 150 µl of RPMI-1640 medium supplemented with 1 mM sodium pyruvate, 0.05 mM 2-mercaptoethanol, 100 U/ml penicillin, and 100 mg/ml streptomycin. Two additional inserts were also prepared with only a monolayer of bEnd.3 cells adding only medium without BMDMs, as a “blank” and as a control monolayer coating the insert. The supernatant from a 72-hour culture of GL-261 glioma cells was added to the lower chambers. After 18 h of incubation at 37°C (5% CO_2_, 95% humidity), the upper part of the inserts was wiped using a cotton swab to remove cells that did not migrate, except for the monolayer control insert, which was not wiped off. Then, the cells on the inserts were stained with the RAL 555 kit (RAL Diagnostics) according to the manufacturer’s instructions. Migrated cells were observed by an Olympus CKX53 light microscope and counted using ImageJ software from 6 images of the central part of each insert. The migration assay was performed in two independent experiments.

### CellTiter-Glo 3D cell viability assay

To determine the effect of BMDMs on the viability of glioma spheroids after 3 and 6 days of co-culture, CellTiter-Glo 3D reagent (Promega) was added and incubated for 30 min in the dark at room temperature. After this time, the relative luminescence units (RLU) were measured using the BioTek Synergy H4 plate reader with Gen5 Software (Agilent Technologies) and an integration time of 0.5 s. The RLU values for the corresponding macrophage populations were subtracted from the RLU values obtained for the co-cultures to obtain the RLU value of glioma spheroids. The CellTiter-Glo 3D cell viability assay was performed in two independent experiments.

### Statistical Analysis

All data were analyzed using GraphPad Prism 10 software (GraphPad Software). The D’Agostino−Pearson omnibus test confirmed the normality of residuals. When the data were consistent with a Gaussian distribution and had equal standard deviation (SD) values, statistical significance was calculated using the parametric one-way ANOVA or the two-way ANOVA followed by Tukey’s multiple comparison post-hoc test. When data were consistent with a Gaussian distribution, but SD values were not equal, the Brown−Forsythe and Welch ANOVA test, followed by Dunnett’s T3 multiple comparisons post-hoc test, was performed. The type of statistical analysis used is described in the captions under the figures. All statistically significant differences are presented in graphs when p<0.05; otherwise, the differences were not significant.

## Results

### Effect of B_4_C nanoparticles on the viability and phenotype of BMDMs

To develop effective cellular carriers, it is crucial to select the nanoparticle concentration that does not affect cell viability, mobility and phenotype. Therefore, in the first stage of the study, the effect of boron carbide nanoparticles on the viability of M0, M1, and M2 bone marrow-derived macrophages after 24 h of exposure was assessed using the MTT assay (Fig. 1A). The results demonstrated boron carbide toxicity toward BMDMs only at higher concentrations above 100 µg/ml, with M1 macrophages being the most sensitive population to B_4_C nanoparticles. The concentration of 100 µg/ml was selected as optimal for further studies, because it did not reduce macrophage viability below 70%.

**Fig. 1.**
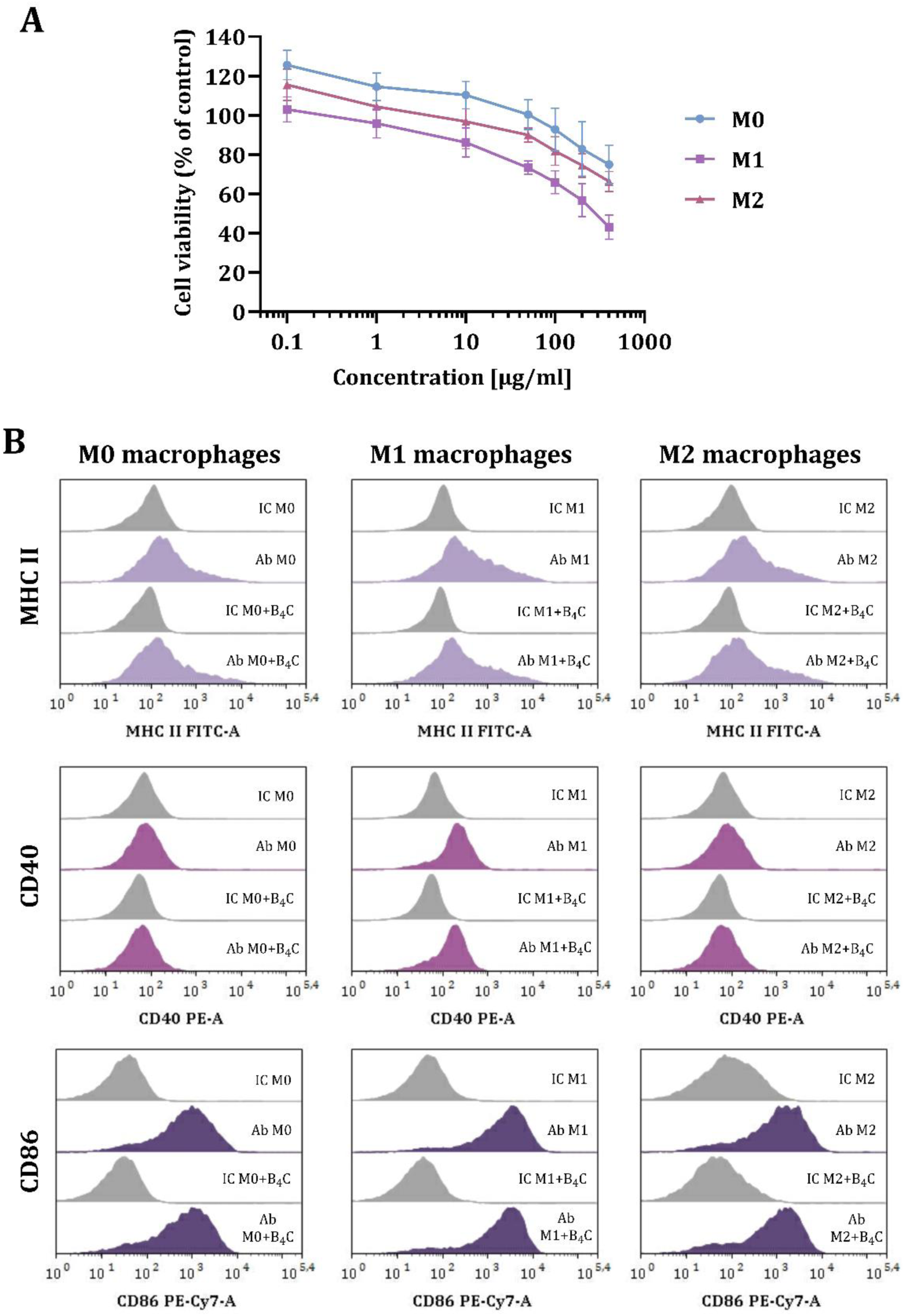
Effect of boron carbide nanoparticles on bone marrow-derived macrophages in three polarization states (M0, M1, M2). **A** MTT viability assay on M0, M1, and M2 macrophages after 24-hour exposure to B_4_C nanoparticles. The graph represents the percentage of viable cells relative to untreated control cells (control = 100%). The results are expressed as the means ± SD calculated for two independent experiments. **B** The phenotypic characterization of M0, M1, and M2 macrophages after 24-hour exposure to B_4_C nanoparticles. Histograms show the mean fluorescence intensity (MFI) of the markers: MHC II, CD40, and CD86 in three populations of BMDMs (F4/80^+^CD11b^+^) with and without B_4_C nanoparticles. The MFI value for each marker after antibody (Ab) staining is compared with the MFI value of the corresponding isotype controls (IC).

Additionally, phenotypic characterization was performed on day 8 of BMDM culture without boron carbide nanoparticles and after 24-hour exposure to 100 µg/ml B_4_C (Fig. 1B). Macrophages were characterized as F4/80 and CD11b positive cells using flow cytometry. Expression of markers such as MHC II, CD40, and CD86 was determined in the BMDM population in three polarization states (M0, M1, and M2). The highest expression of all antigens was observed in M1 macrophages. Furthermore, the obtained results showed that boron carbide nanoparticles had no significant effect on the mean fluorescence intensity (MFI) of all markers. This confirms the stability of the macrophage phenotype after 24-hour incubation with B_4_C nanoparticles at a selected concentration.

### Uptake and accumulation of B_4_C nanoparticles in BMDMs

In the next stage, the uptake of boron carbide nanoparticles by M0, M1, and M2 bone marrow-derived macrophages was compared. For this purpose, BMDMs were incubated for 24 h with B_4_C nanoparticles at a concentration of 100 µg/ml. Initially, macrophages with accumulated nanoparticles were visualized using holotomographic microscopy (Fig. 2). Three-dimensional (3D) images were generated using a pseudo-coloring technique based on differences in refractive index (RI). In the 3D visualizations, green color indicates cell components (organelles, cytoplasm) that exhibit a lower RI (<1.4) than the boron carbide nanoparticles (RI>1.4), which are marked in red. A high intracellular accumulation of B_4_C nanoparticles was observed in all macrophage populations, but the highest number of nanoparticles was identified in M1 macrophages.

**Fig. 2.**
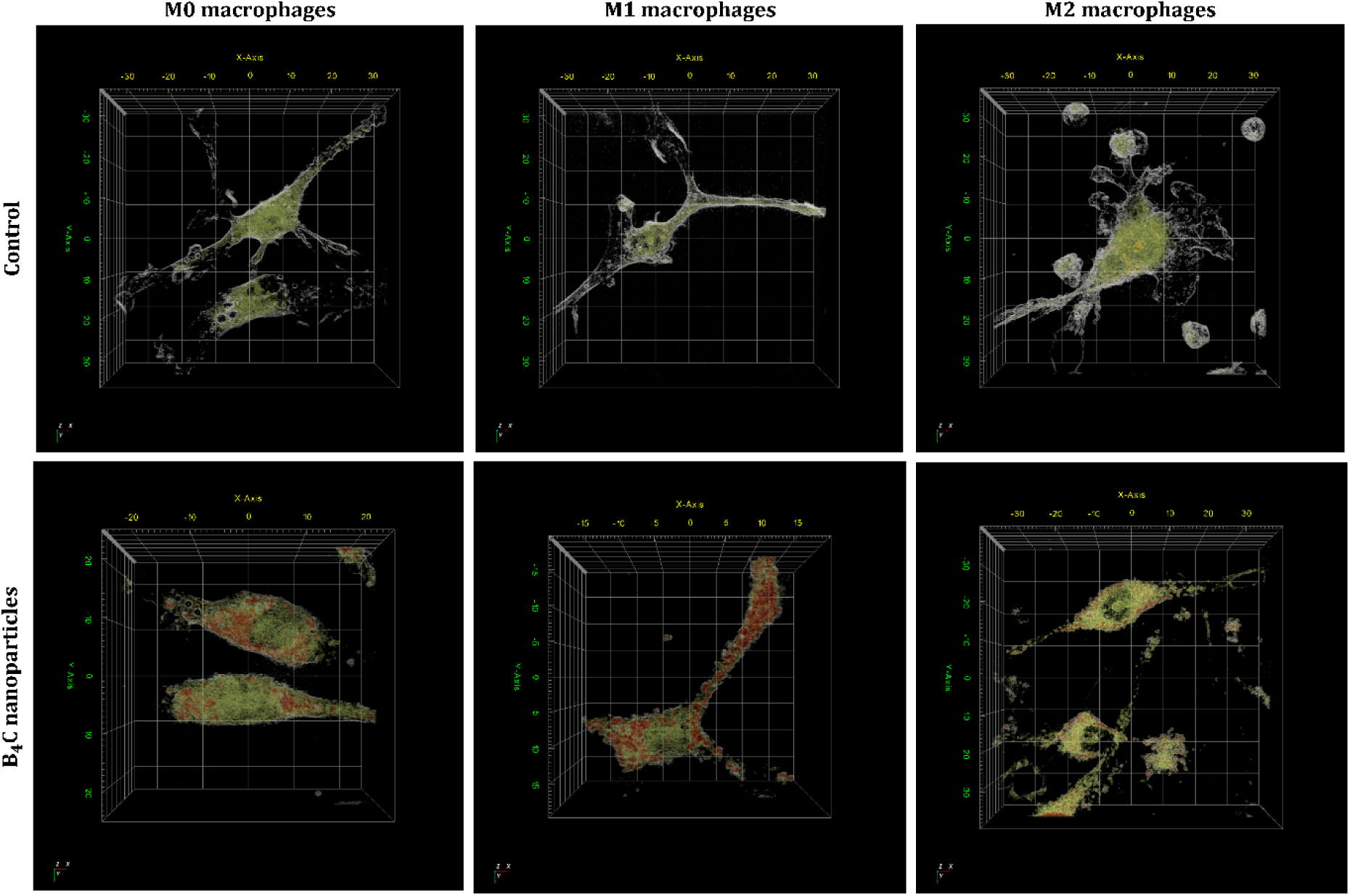
Three-dimensional holotomographic visualization of M0, M1, and M2 bone marrow-derived macrophages with engulfed boron carbide nanoparticles compared to untreated cells, using pseudo-coloring based on refractive index (RI) ranges. Green structures have an RI of up to 1.4 and mark the interior of the cells (cytoplasm and organelles), while the red color corresponds to boron carbide nanoparticles with an RI above 1.4.

After an initial qualitative assessment of uptake, inductively coupled plasma mass spectrometry (ICP-MS) was used to quantitatively analyze the boron concentration engulfed by BMDMs (Fig. 3). The analysis showed statistically significant differences in boron concentrations in all macrophage populations after incubation with B_4_C nanoparticles compared to untreated cells, where the boron content was negligible (<0.35 mg/l per 10^6^ cells). The highest boron concentration was detected in M1 macrophages (21.53 ± 1.64 mg/l per 10^6^ cells), while the concentrations in M0 and M2 macrophages were 14.27 ± 1.36 mg/l and 16.72 ± 1.93 mg/l per 10^6^ cells, respectively. The highest accumulation of boron carbide nanoparticles in M1 macrophages was associated with the greatest observed decrease in the viability of these cells among all BMDM populations.

**Fig. 3.**
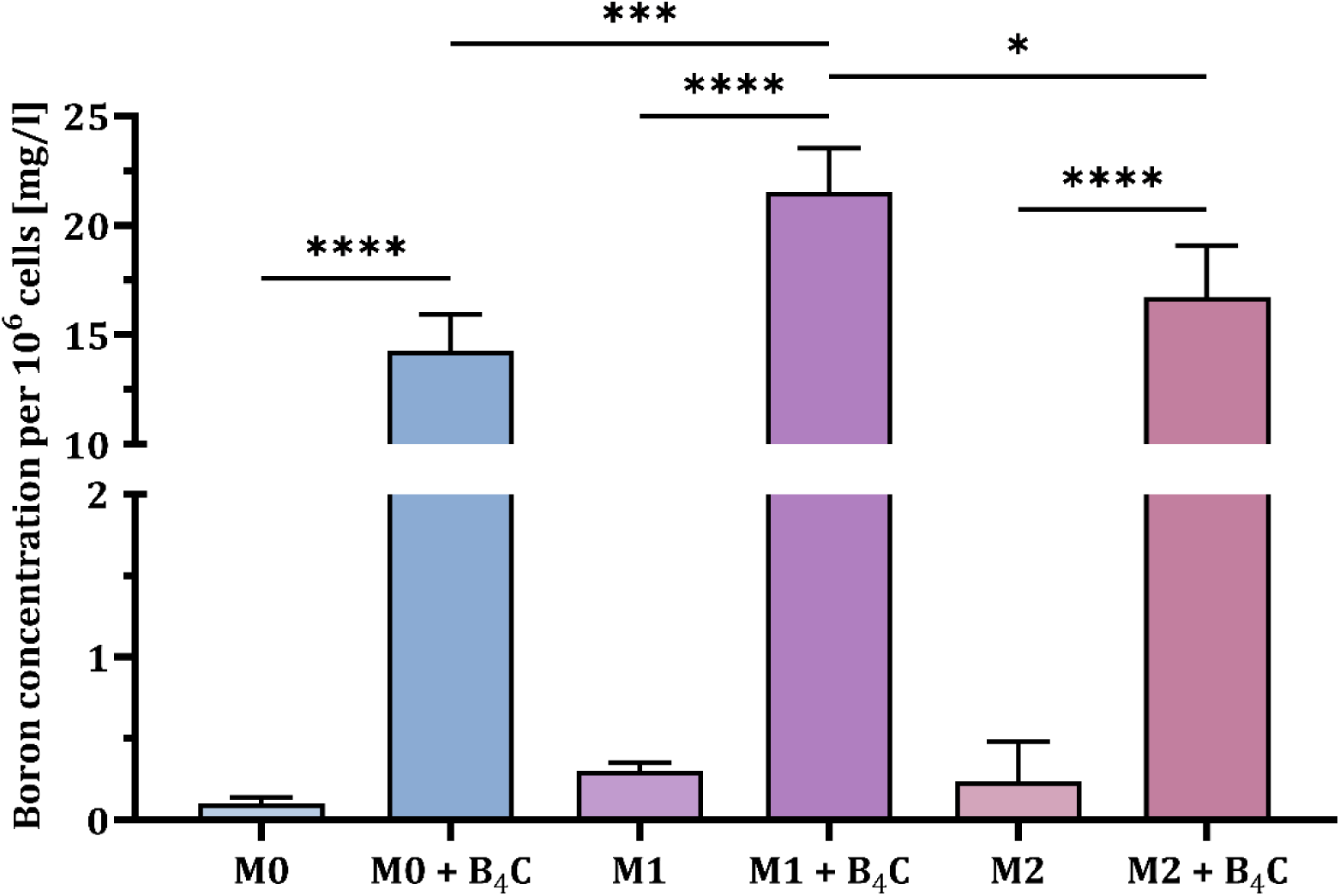
Boron concentration per 10^6^ bone marrow-derived macrophages in three populations (M0, M1, and M2) after 24 h of incubation with boron carbide nanoparticles compared to untreated control cells. The results are expressed as the means + SD calculated from three independent biological samples. The differences between groups were calculated using the one-way ANOVA followed by Tukey’s multiple comparison post-hoc test (*p < 0.05; ***p < 0.001; ****p < 0.0001).

### Migration ability of BMDMs

Another key aspect of effective cellular carriers is their ability to deliver engulfed nanoparticles to the target site. Therefore, the spontaneous migration ability of three macrophage populations (M0, M1, and M2) with and without accumulated boron carbide nanoparticles was compared using the scratch assay (Fig. 4). The progress of cell migration was observed under a light microscope after 4 and 24 h. The results showed that M0 macrophages migrated faster with B_4_C nanoparticles than without them. In contrast, M1 macrophages with nanoparticles migrated more slowly than those without nanoparticles. The migration rate of M2 macrophages was the same regardless of the presence of nanoparticles. These observations indicate that efficient B_4_C uptake by M1 macrophages affects not only their viability but also their mobility.

**Fig. 4.**
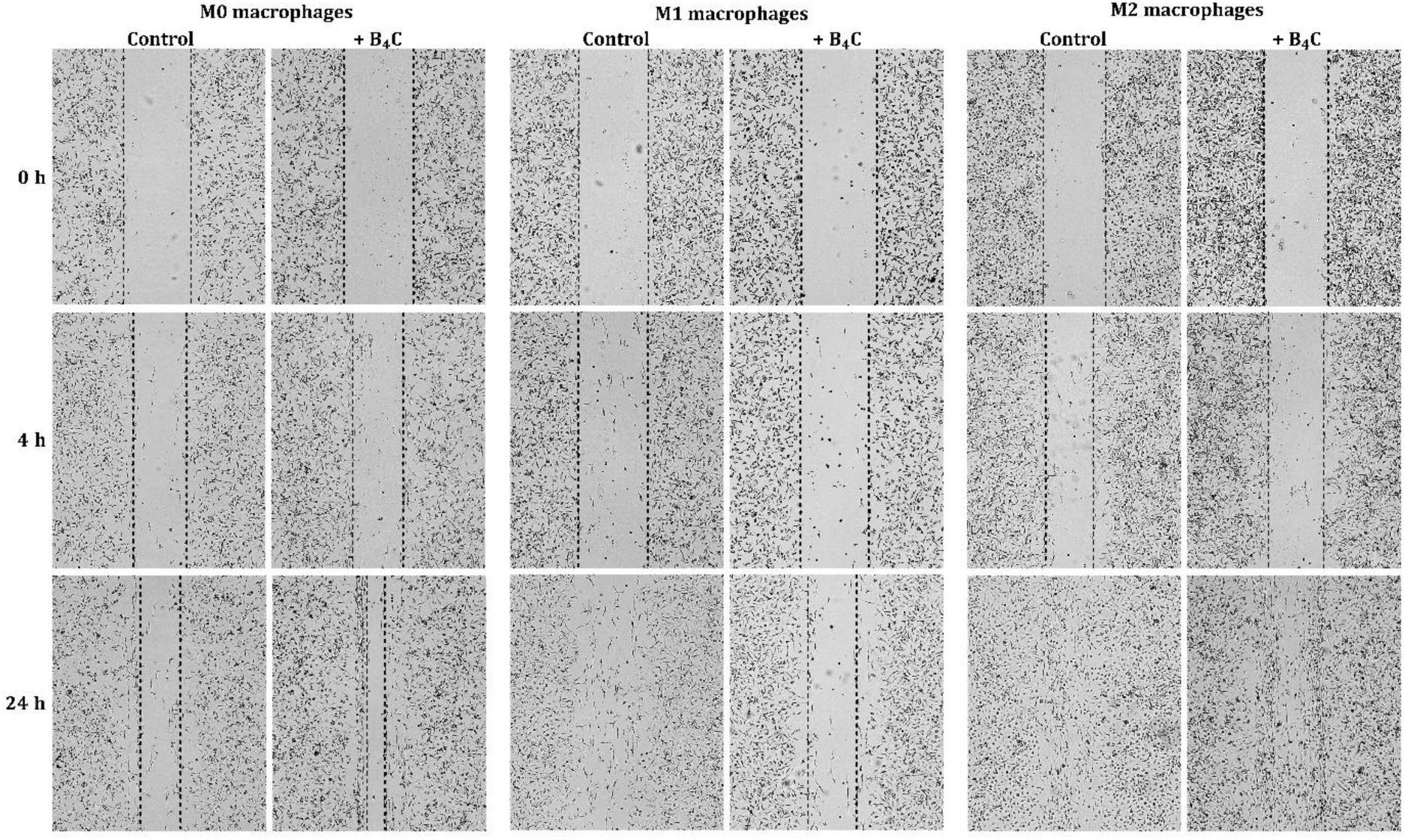
Assessment of the spontaneous migration ability of M0, M1, and M2 bone marrow-derived macrophages with and without engulfed boron carbide nanoparticles based on the scratch assay after 4 and 24 h.

### Crossing the blood-brain barrier by BMDMs

In the case of brain tumors, the ability of the cellular carriers to cross the blood-brain barrier is crucial. To compare the ability of three BMDM populations with and without accumulated boron carbide nanoparticles to migrate across the BBB toward the supernatant of GL-261 glioma cells, a Transwell migration assay was performed. First, the concentration of the chemokine CCL2 in the supernatants from 24-, 48-, and 72-hour cultures of GL-261 cells was assessed by ELISA due to its important role in macrophage recruitment to the TME (Fig. 5A). The results showed the highest concentration of CCL2 in 72-hour culture supernatants of GL-261 cells at approximately 112 pg/ml and it was selected as a chemoattractant for the Transwell migration assay.

**Fig. 5.**
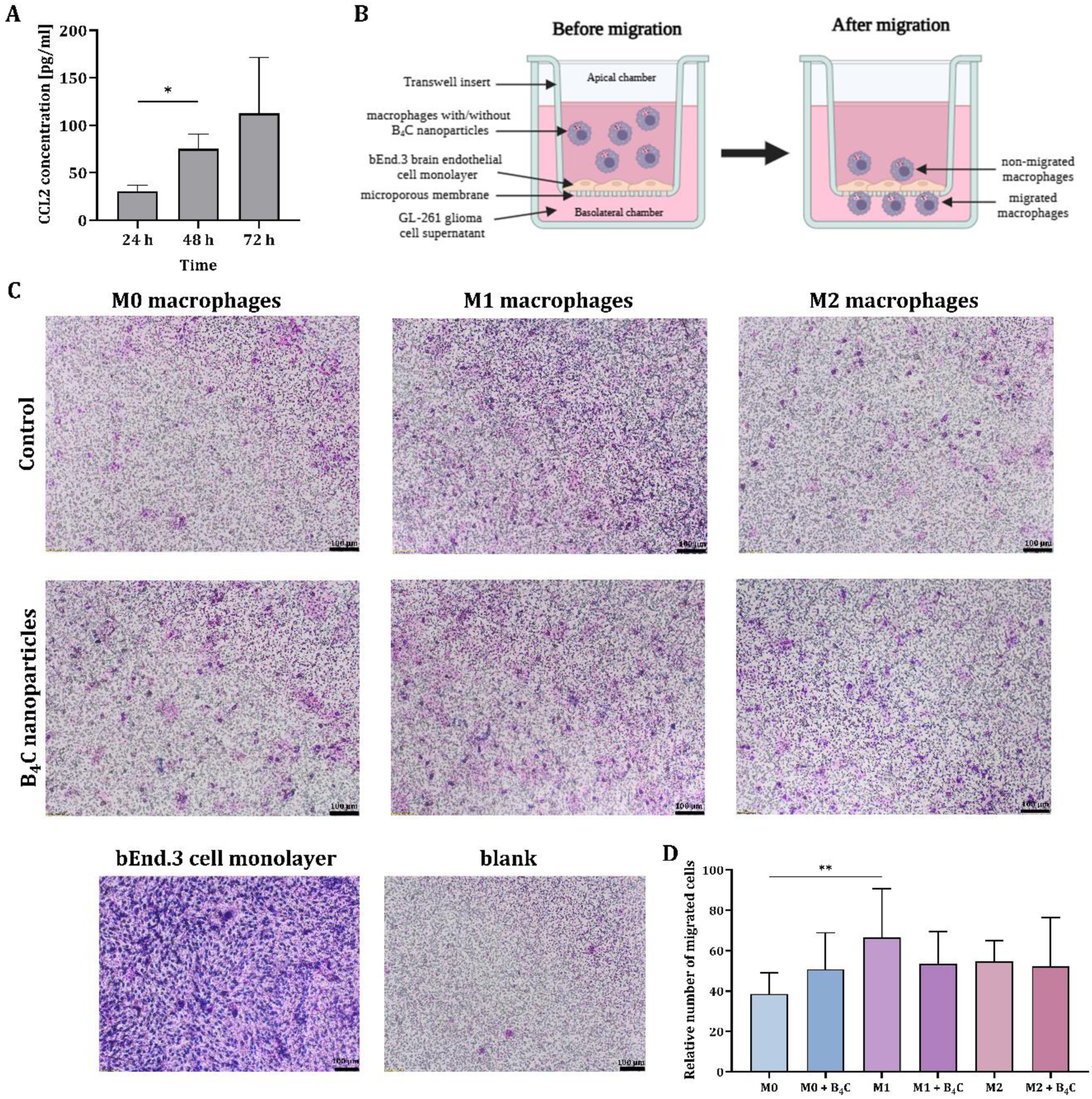
**A** CCL2 concentrations in 24, 48, and 72-hour supernatant from GL-261 glioma cells determined by ELISA. **B** Schematic representation of the Transwell migration assay. **C** Images of migrated M0, M1, and M2 bone marrow-derived macrophages with and without accumulated boron carbide nanoparticles through a bEnd.3 brain endothelial cell monolayer toward 72-hour supernatant from GL-261 glioma cells containing CCL2. **D** Comparison of the average number of migrated cells counted from 6 images of the central part of each insert. The results are expressed as the means + SD calculated from two independent experiments. Differences between groups were calculated using (A) the Brown−Forsythe and Welch ANOVA test, followed by Dunnett’s T3 multiple comparisons post-hoc test (*p < 0.05), (D) one-way ANOVA followed by Tukey’s multiple comparison post-hoc test (**p < 0.01).

To create an *in vitro* model of the blood-brain barrier, bEnd.3 brain endothelial cells were used to coat Transwell inserts. Then, M0, M1, and M2 bone marrow-derived macrophages, both with and without accumulated boron carbide nanoparticles, were applied to an insert coated with endothelial cells (apical chamber). The wells under the inserts (basolateral chamber) were filled with 72-hour supernatant from GL-261 glioma cells (Fig. 5B). Inserts coated with endothelial cells without added BMDMs served as a control for bEnd.3 cell migration (blank) and as a control for monolayer integrity.

After 18 h of migration, the migrated macrophages on the inserts were stained. The observations obtained in the Transwell test were similar to those in the scratch assay. The number of migrated M0 macrophages with B_4_C nanoparticles was greater than that of M0 macrophages without nanoparticles (Fig. 5C, D). M1 macrophages without B_4_C nanoparticles migrated better than with nanoparticles. For M2 macrophages, both with and without nanoparticles, the number of migrated cells was similar. Despite the observed differences within the population, all types of macrophages (M0, M1, and M2) loaded with nanoparticles migrated comparably.

### Effect of BMDMs on the viability of GL-261 spheroids and changes in CCL2 production in co-culture

The proven ability of bone marrow-derived macrophages loaded with B_4_C nanoparticles to cross the brain endothelial cell monolayer, mimicking the BBB, constitutes their great advantage as cellular carriers. Therefore, subsequent steps were focused on assessing possible interactions between macrophages and tumor cells using 3D cultures. For this purpose, 10-day-old GL-261 spheroids were co-cultured with three BMDM populations (M0, M1, and M2), with or without accumulated boron carbide nanoparticles. Control spheroid culture formed from GL-261 glioma cells and co-culture with Calcein AM Green-labeled macrophages were imaged using a fluorescence microscope (Fig. 6A). Co-culture of spheroids and BMDMs was conducted for 3 and 6 days. After this time, the effect of macrophages on spheroid viability was assessed using the CellTiter-Glo assay, in which relative luminescence units (RLU) correspond to cell viability (Fig. 6B). After 3 days, a slight increase in the viability of glioma spheroids was observed in co-cultures with macrophages loaded or unloaded with B_4_C nanoparticles compared to control spheroids. However, after 6 days of co-culture, a differential survival of tumor cells was demonstrated. Importantly, macrophages loaded with nanoparticles did not affect spheroid viability, whereas contact with control macrophages significantly increased the survival of glioma cells.

**Fig. 6.**
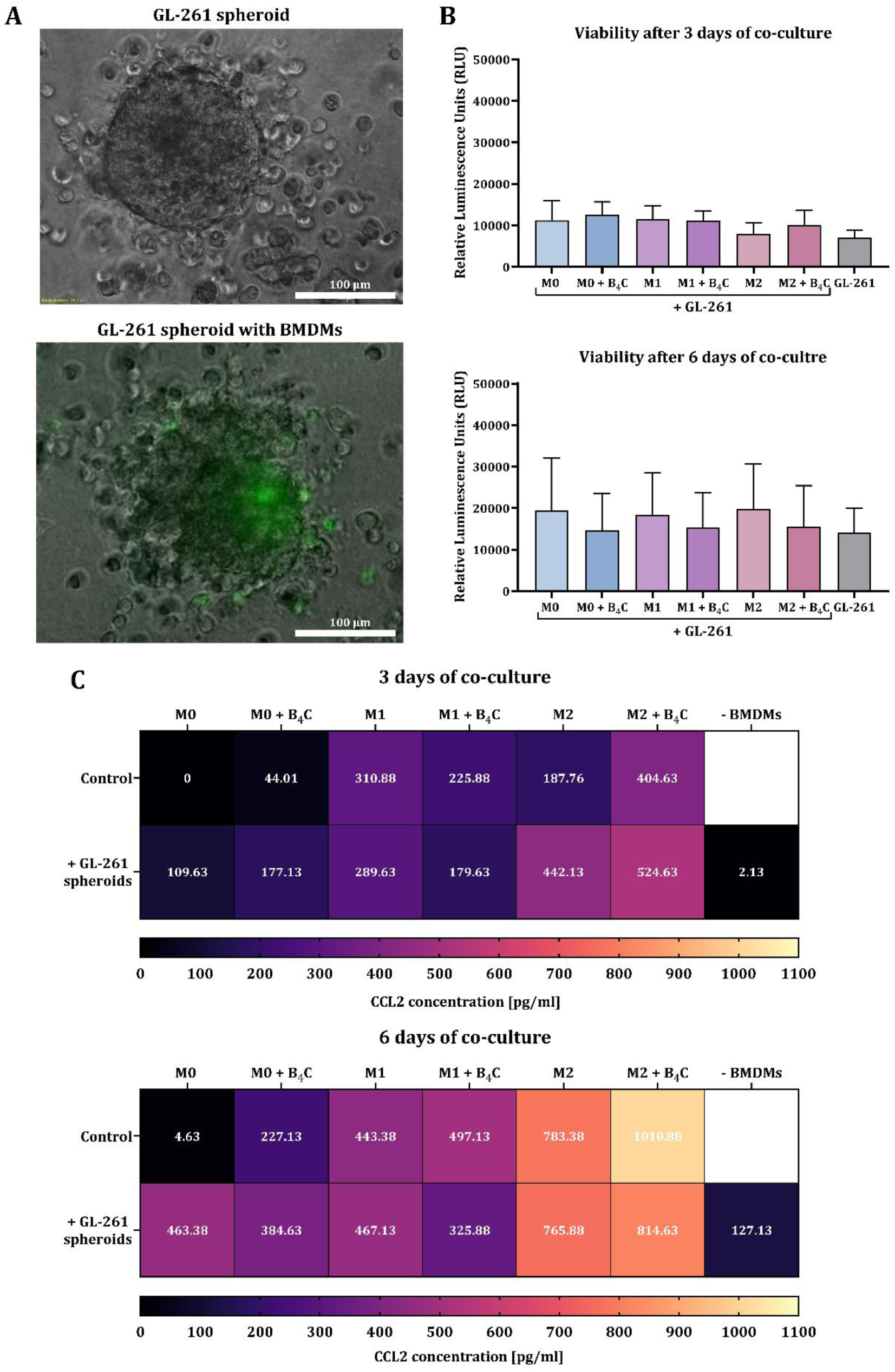
Effect of M0, M1, and M2 bone marrow-derived macrophages loaded with or without boron carbide nanoparticles on the spheroids obtained from GL-261 glioma cells after 3 and 6 days of co-culture. **A** Brightfield image of a spheroid formed from GL-261 glioma cells. Green fluorescence indicates BMDMs labeled with Calcein AM Green. **B** Viability determined by CellTiter-Glo 3D assay after 3 and 6 days of co-culture. The relative luminescence units (RLU) are derived from GL-261 spheroids after subtracting the RLU value for the corresponding control macrophage population. The results are expressed as the means + SD calculated from two independent experiments. **C** Concentration of CCL2 in co-culture of glioma spheroids with BMDMs after 3 and 6 days, compared to the production by macrophages and spheroids alone.

Additionally, CCL2 levels were assessed in co-cultures of BMDMs and GL-261 spheroids after 3 and 6 days using ELISA (Fig. 6C). The results showed higher CCL2 production after 6 days than after 3 days, both in co-culture and in the culture of macrophages and spheroids alone. After 3 days of culture, M1 macrophages loaded with B_4_C nanoparticles produced less CCL2 than M1 without nanoparticles. The same tendency was also observed in co-culture with spheroids. However, in M0 and M2 populations, the effect was reversed. After 6 days of culture, M0, M1, and M2 macrophages loaded with B_4_C nanoparticles produced more CCL2 than those without nanoparticles. However, when spheroids were co-cultured with nanoparticle-loaded M0 and M1 macrophages, CCL2 levels were lower than in co-cultures with BMDMs without engulfed nanoparticles. M2 macrophages with and without accumulated nanoparticles produced the highest levels of CCL2 among all populations, which was also observed in the CCL2 levels in co-culture with spheroids.

### Effect of GL-261 spheroids on changes in the phenotype of BMDMs

After 3 and 6 days of co-culture, the effect of GL-261 spheroids on changes in macrophage marker expression was assessed. For this purpose, the phenotypic characterization of three BMDM populations with and without accumulated boron carbide nanoparticles was performed using flow cytometry (Fig. 7). The expression of MHC II and CD40 on the surface of M0, M1, and M2 macrophages increased after contact with spheroids at both time points, with the highest levels of these molecules found in the M1 population. However, increases in marker expression were greater in all unloaded macrophage populations than in nanoparticle-loaded cells. No statistically significant differences in MHC II and CD40 surface expression between days 3 and 6 of culture were observed.

**Fig. 7.**
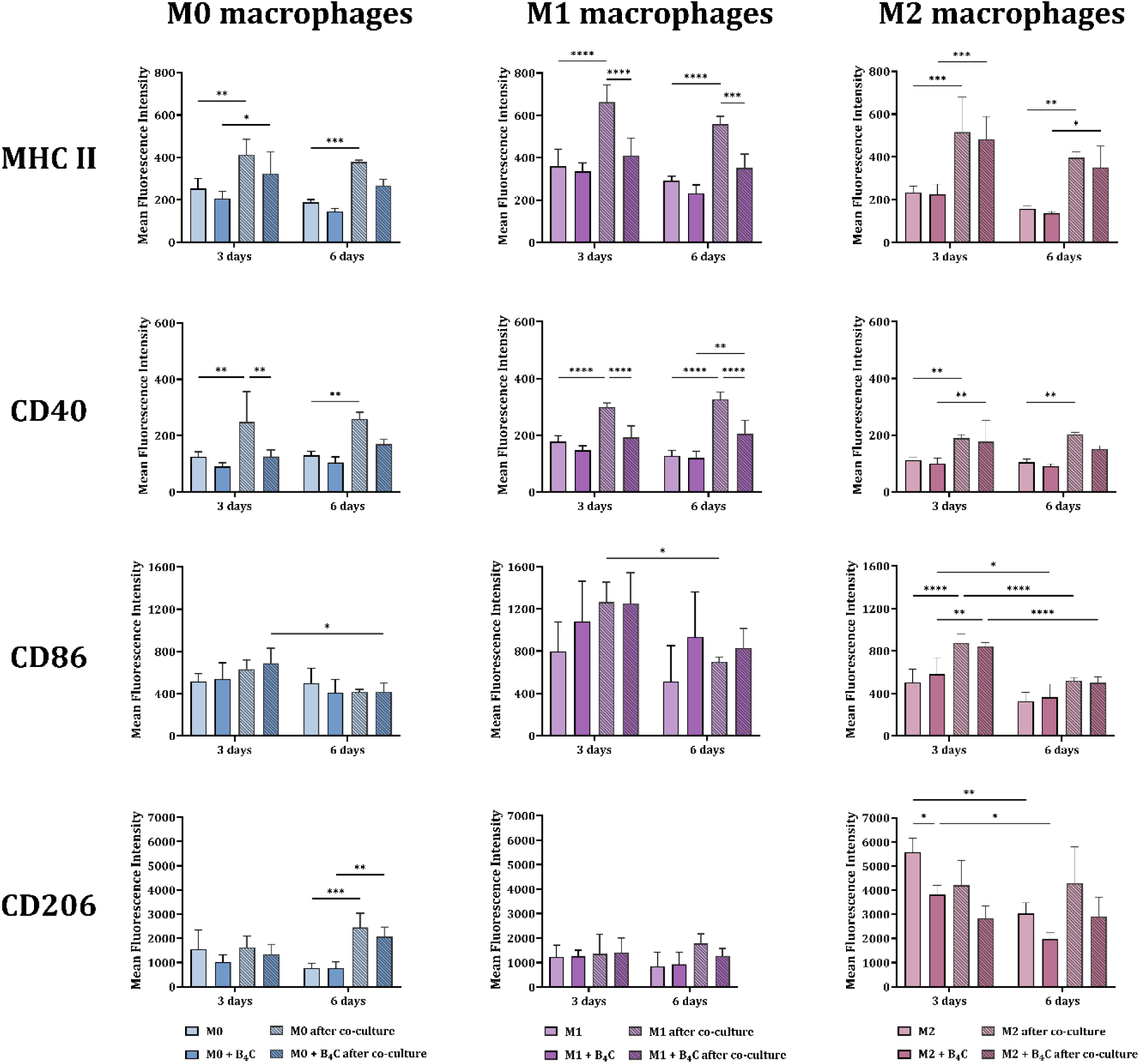
Phenotypic characterization of bone marrow-derived macrophages loaded with or without boron carbide nanoparticles after 3 and 6 days of co-culture with GL-261 glioma spheroids. The mean fluorescence intensity of markers: MHC II, CD40, CD86, and CD206 was determined for the BMDM population (F4/80^+^CD11b^+^). The results are expressed as the means + SD calculated from two independent experiments. Differences between groups were calculated using two-way ANOVA followed by Tukey’s multiple comparison post-hoc test (*p < 0.05; **p < 0.01; ***p < 0.001; ****p < 0.0001).

At the second time point, both in culture with and without spheroids, a decrease in CD86 expression was observed in all macrophage populations. However, regardless of B_4_C loading, M1 macrophages showed the highest expression of this molecule compared to the other populations.

After 6 days of culture, differences in mannose receptor expression levels were observed. In M0 macrophages, both with and without nanoparticles, an increase in CD206 expression was noted compared to control M0 macrophages. In M2 macrophages loaded with B_4_C nanoparticles, CD206 expression was lower compared to M2 macrophages without nanoparticles. Meanwhile, the presence of tumor spheroids did not cause repolarization of M1 macrophages towards the M2 phenotype, as evidenced by the stable expression of CD206 on their surface not only on day 6 but also on day 3 of co-culture.

## Discussion

The use of macrophages as cellular carriers in boron neutron capture therapy is an original and innovative strategy. In our previous work [16], we demonstrated the potential of macrophages originating from a cell line and bone marrow-derived macrophages to accumulate two boron carbide preparations differing in nanoparticle size. To continue and extend our research, we selected smaller nanoparticles (~32 nm), which showed less toxicity to macrophages than larger ones (~80 nm). Furthermore, we focused on bone marrow-derived macrophages, which, as primary cells, better mimic *in vivo* macrophage physiology than those derived from cell lines.

To ensure efficient migration of macrophages to the target site, it is important to minimize the impact of engulfed compounds on their viability, phenotype, and normal function. Therefore, based on the MTT viability assay, we selected a B_4_C nanoparticle concentration of 100 μg/ml for further studies, which provides a high level of boron without significant toxicity during 24-hour incubation with BMDMs (Fig. 1A). We also confirmed that this concentration did not influence changes in the expression of MHC II, CD40, and CD86 molecules on the surface of all macrophage populations after 24 h of exposure (Fig. 1B). Moreover, in previous studies, we demonstrated no changes in the production of cytokines such as IL-1β, IL-6, IL-10, and TNF-α by all macrophage populations after incubation with B_4_C nanoparticles at a concentration of 100 μg/ml [16].

Another key aspect for obtaining cellular carriers is the effective internalization of the compound by macrophages. Therefore, we proved the uptake of boron carbide nanoparticles by M0, M1, and M2 macrophages over a 24-hour incubation period. Holotomographic microscopy visualization and ICP-MS analysis confirmed significant accumulation of nanoparticles in all macrophage populations (Figs. 2 and 3). The highest boron concentration was detected in M1 macrophages. This may be due to the greater ability of activated M1 macrophages to phagocytize large amounts of particles compared to M0 and M2 macrophages. Moreover, it is related to the natural function of M1 macrophages in the immune response, which involves phagocytizing pathogens and presenting antigens to activate T helper 1 cells [20]. The differential ability of M0, M1, and M2 macrophages to accumulate nanoparticles observed in our study corresponds to other published results obtained for murine bone marrow-derived macrophages. In a study by Pang et al., bone marrow-derived M1 macrophages demonstrated a significantly higher capacity to engulf doxorubicin-loaded poly(lactide-co-glycolide) (PLGA) nanoparticles compared to M0 macrophages [21]. Similarly, Qie et al. showed that M1 macrophages had the highest ability to phagocytose carboxylic acid-terminated fluorescently labeled polystyrene nanoparticles compared to M0 and M2 macrophages [22]. Furthermore, Müller et al. proved that M1 macrophages had higher phagocytic activity toward rabbit IgG-coated latex particles than M0 and M2 macrophages [23].

The next important step in the development of cellular carriers was to evaluate the ability of macrophages loaded with boron carbide nanoparticles to spontaneously motility and migrate toward the tumor microenvironment. Based on the scratch assay, it was confirmed that among all macrophage populations with engulfed B_4_C nanoparticles, M2 macrophages are characterized by the highest mobility (Fig. 4). In contrast, M1 macrophages loaded with nanoparticles migrated the slowest. Similarly, in the work of Li et al., based on the calculated random motility coefficient, M2 bone marrow-derived macrophages showed the highest migration ability compared to M0 and M1 macrophages [24]. This commonly observed phenomenon is related to stronger adhesion properties of M1 macrophages, which reduces their overall motility, while M2 macrophages have a more dynamic and less adherent migration profile [25]. This is due to the physiological functions of these phenotypically distinct macrophage populations. M2 macrophages play a crucial role in relieving inflammation and initiating the tissue repair process, in which migration to sites of injury is essential. The strong adhesion and limited motility of M1 macrophages allow them to accumulate at the site of inflammation, release inflammatory mediators, and eliminate pathogens [26]. Moreover, in this work, the slowest migration of M1 macrophages may be associated with the most efficient uptake of B_4_C nanoparticles, as confirmed by the ICP-MS results.

The natural ability of macrophages to cross the blood-brain barrier makes them extremely attractive cellular carriers for brain tumor therapies. However, to demonstrate that bone marrow-derived macrophages loaded with B_4_C nanoparticles can cross the BBB, we used the Transwell system. To create an *in vitro* model of the BBB, we coated Transwell inserts with bEnd.3 brain endothelial cells, which form tight junctions with strong intercellular barrier properties [27]. As a chemoattractant, we utilized the 72-hour supernatant from GL-261 glioma cells, which was rich in CCL2 (Fig. 5A). The Transwell assay results correlated with the scratch assay observations. In the case of M0 macrophages, B_4_C stimulated migration, whereas in M1 macrophages, it slightly inhibited migration compared to control macrophages (Figs. 5C and D). A similar phenomenon was observed by Pang et al., who showed that doxorubicin-loaded PLGA nanoparticles reduced the migratory ability of M1 bone marrow-derived macrophages through a monolayer of human umbilical vein endothelial cells (HUVECs) toward human U87 glioma cells in a Transwell model [21]. Nonetheless, our results confirmed that all macrophage populations (M0, M1, and M2) loaded with boron carbide nanoparticles showed a similar ability to migrate through the endothelial cell monolayer toward the supernatant of GL-261 cells.

To assess the further fate of macrophages after crossing the blood-brain barrier, interactions between macrophages and glioma spheroids, reflecting the tumor microenvironment, were analyzed. After 3 days of co-culture, all macrophage populations with and without nanoparticles slightly increased the viability of glioma spheroids (Fig. 6B). However, after 6 days, macrophages with accumulated B_4_C showed no effect on tumor cell survival. While control BMDMs significantly increased spheroid viability. The ability of macrophages in different polarization states (M0, M1, and M2) to stimulate the growth of tumor spheroids was also described by Francois et al. [28]. The lack of increased survival of glioma cells under the influence of nanoparticle-loaded macrophages is a very beneficial phenomenon for the effectiveness of the proposed therapeutic strategy.

Considering the crucial role of CCL2 in the recruitment of monocytes and macrophages to the TME, its concentration was assessed in co-cultures of macrophages with glioma spheroids. The observed significant increase in CCL2 production in all macrophage-spheroid co-cultures after 6 days compared to day 3 may not only act as a positive feedback loop for recruiting further macrophage carriers but also unwanted cells such as Tregs and MDSCs, promoting tumor progression (Fig. 6C). Moreover, CCL2 has been shown to promote macrophage polarization into the M2 phenotype and an immunosuppressive tumor environment [29]. Importantly, our results showed that 3- and 6-day co-cultures of spheroids with M1 macrophages loaded with B_4_C nanoparticles resulted in lower CCL2 levels compared to co-cultures with control M1 macrophages and other macrophage populations. Further analysis of the macrophage phenotype after co-culture with spheroids revealed that M1 macrophages did not increase CD206 expression (Fig. 7). Moreover, M2 macrophages with B_4_C nanoparticles showed a decrease in CD206 expression after co-culture compared to control macrophages. This suggests that boron carbide may promote the repolarization of M2 macrophages into M1 macrophages, but also prevent the negative effects of the TME on M1 macrophages, and support their antitumor phenotype. The ability of some nanoparticles to repolarize M2 macrophages into the M1 phenotype has already been described. For example, such repolarization induced by iron oxide nanoparticles resulted in inhibition of tumor growth in mice [30]. Moreover, the observed increased expression of markers characteristic for the M1 phenotype (MHC II, CD40, and CD86) in all BMDM populations after co-culture may indicate the activation of macrophages toward immunostimulation and the potential ability to present antigens and activate T lymphocytes. These observations are supported by studies showing that spheroids obtained from certain cell lines, such as MCF-7, can polarize macrophages more strongly toward the M1 than the M2 phenotype [31]. A more common phenomenon is the repolarization of M1 macrophages toward the M2 phenotype caused by the immunosuppressive effects of the tumor microenvironment. However, the ability of M1 macrophages with engulfed B_4_C nanoparticles to maintain this phenotype in the TME is highly desirable for supporting anticancer therapy.

## Conclusions

Our studies indicate that bone marrow-derived macrophages are promising carriers of boron carbide for BNCT. They efficiently engulf B_4_C nanoparticles and migrate across the BBB toward the tumor site. M1 macrophages appear to be the best candidate for carriers because they not only engulf nanoparticles most efficiently but also, as a result, do not repolarize toward M2 phenotype and produce less CCL2 compared to other populations. Furthermore, their anticancer properties may further support the eradication of tumor cells. Considering the specific role of macrophages in the immune response and the originality of the mentioned therapeutic strategy, it may be a new type of radioimmunotherapy.

## Author contributions

AR, BSO and EPP contributed to the conceptualization, writing - original draft; AR, BSO, MCM, PR, KW contributed to the methodology; AR, BSO, AS, KWC, JM, DK, PŻ, MCM, PR, KW, ZP, EPP contributed to the investigation and writing – review and editing; BSO and EPP contributed to the supervision; AR contributed to the visualization, formal analysis, data curation, project administration, funding acquisition and resources. All authors read and approved the final manuscript.

## Abbreviations

3D: Three-dimensional
4He: Helium-4 isotope - alpha particles
7Li: Lithium-7 isotope
10B: Boron-10 isotope
B_4_C: Boron carbide
BBB: Blood-brain barrier
BMDMs: Bone marrow-derived macrophages
BNCT: Boron neutron capture therapy
CCL2: C-C motif chemokine ligand 2
CCR2: C-C chemokine receptor type 2
DLS: Dynamic light scattering
DMEM: Dulbecco’s modified Eagle’s medium
EGF: Epidermal growth factor
FBS: Fetal bovine serum
FGF: Fibroblast growth factor
ICP-MS: Inductively coupled plasma mass spectrometry
IFN-γ: Interferon-γ
IL: Interleukin
LDV: Laser Doppler velocimetry
LET: High linear energy transfer
LPS: Lipopolysaccharide
MCP-1: Monocyte chemotactic protein-1
M-CSF: Macrophage colony-stimulating factor
MDSCs: Myeloid-derived suppressor cells
MFI: Mean fluorescence intensity
MHC II: Major histocompatibility complex class II
MTT: 3-(4,5-dimethylthiazol-2-yl)-2,5-diphenyltetrazolium bromide
PLGA: Poly(lactide-co-glycolide
RI: Refractive index
RLU: Relative luminescence units
SEM: Scanning electron microscope
TAMs: Tumor-associated macrophages
TEM: Transmission electron microscopy
TME: Tumor microenvironment
TNF-α: Tumor necrosis factor α
Tregs: Regulatory T lymphocytes

